# Determining preclinical safety of Aclarubicin in pediatric malignancies

**DOI:** 10.1101/2025.05.20.654130

**Authors:** Darleen S Tu, Aaron K Olson, Kimberly S Waggie, Nicolas M Garcia, Virginia J Hoglund, Stephanie Walter, Jenna R Rosinski, Harini Sadeeshkumar, Radhika Patel, Erolcan Sayar, Michael C Haffner, Lisa Maves, Jacques Neefjes, Jay F Sarthy, Elizabeth R Lawlor, Shireen S Ganapathi

## Abstract

**Background:** Anthracyclines are among the most effective chemotherapeutic agents used to treat pediatric malignancies. However, their clinical use is limited by dose-dependent toxicities, particularly cardiotoxicity and secondary malignancies. Aclarubicin (Acla) is an anthracycline derivative that induces chromatin damage while sparing DNA, offering potential therapeutic benefit with reduced longterm toxicity.

**Methods:** We evaluated the anti-tumor efficacy and safety profile of Acla in multiple *in vitro* pediatric cancer models and *in vivo* mouse models designed to mimic anthracycline re-treatment following prior doxorubicin (Doxo) exposure. Tumor growth, genotoxic stress, survival, and organ toxicity were assessed.

**Results:** Acla demonstrated robust anti-tumor activity comparable to Doxo across diverse pediatric *in vitro* models. Unlike Doxo, Acla treatment did not induce significant genotoxic stress. *In vivo*, mice receiving Acla after Doxo exposure showed no evidence of cumulative cardiotoxicity or end-organ damage. In contrast, a second course of Doxo led to significant toxic mortality, but was surprisingly not attributable to classic cardiac injury.

**Conclusion:** Our study highlights Acla as a promising anthracycline derivative for pediatric cancers, with potent anti-tumor efficacy and a superior safety profile, even following prior anthracycline exposure. These results support continued investigation of chromatin-damaging anthracyclines that can kill pediatric cancer cells without inducing genotoxic stress. In addition, our studies underscore the need to refine preclinical models to better understand both acute and chronic anthracycline toxicities in pediatric and adolescent populations.

## Introduction

Anthracyclines remain one of the most important class of chemotherapeutic agents used to treat pediatric and adolescent and young adult (AYA) malignancies. Nevertheless, while dose escalation of anthracyclines has improved survival, it is limited by dose-dependent toxicities including infertility, secondary malignancies, and cardiotoxicity(1–4). These dose-dependent toxicities on healthy tissue limit further use in upfront therapy or at relapse(4).

Anthracyclines function as topoisomerase II (topo II) poisons, inducing double-strand DNA breaks (dsDNA) and apoptosis(5). Its ability to poison topo II leads to generation of free radicals, and targeted Topo IIβ poisoning in cardiomyocytes induces cardiac toxicity(4, 6, 7). However, it was recently discovered that anthracyclines also cause chromatin damage by evicting histones leading to epigenetic and transcriptional alterations and subsequent tumor cell death(8–11). Unlike etoposide, which solely causes DNA damage(8), anthracyclines attenuate DNA repair through H2AX phosphorylation. Importantly, chromatin and DNA damage occur at separate genomic loci, indicating independent mechanisms(12). Emerging evidence suggests that chromatin damage contributes to antitumor efficacy without driving off-target toxicity(12, 13).

Acla is a chromatin-damaging anthracycline with minimal genotoxicity due to its distinct structure (12–14). Acla has been used for decades in Europe, Japan, and China to treat elderly patients with acute myeloid leukemia (AML) due to its favorable toxicity profile(15–17). Recent preclinical work demonstrated equivalent anti-tumor efficacy of Acla and Doxo, while only high doses of Doxo inducing significant secondary off target toxicities-cardiotoxicity and hematologic secondary malignancies(12, 13). The biodistribution of Acla is different from Doxo, and Acla preferentially accumulates in hematopoietic organs such as thymus, spleen and bone marrow(12). With increasing recognition of chromatin-mediated anti-cancer effects in anthracyclines, Acla offers a promising strategy to preserve therapeutic efficacy while reducing long-term toxicities in pediatric and AYA malignancies.

Here we evaluated the preclinical efficacy of Acla and Doxo across a panel of pediatric cancer cell lines and found comparable *in vitro* cytotoxicity, independent of genotoxic damage. Given that patients with relapsed disease often approach lifetime anthracycline limits, we modeled Acla administration following prior Doxo exposure. While additional Doxo led to significant toxicity and premature mortality, Acla was well tolerated in pretreated C57BL/6J mice and did not induce significant organ dysfunction. These findings support the anti-tumor efficacy of Acla, and the potential and feasibility of Acla in the relapsed pediatric cancer setting.

## Materials and Methods

### Cell lines

CHLA10-Ewing sarcoma (EwS), L-428-Hodgkin Lymphoma (HL), and Raji-Burkitt Lymphoma (BL) cell lines were obtained from ATCC and COG cell line repositories (https://www.childrensoncologygroup.org/). CHLA10 was cultured in IMDM (Fisher) supplemented with 1% L-glutamine and 20% fetal bovine serum (FBS) and 1X Insulin-Transferrin-Selenium-Ethanolamine (Gibco). Raji and L-428 were cultured in RPMI 1640 (Gibco) with 10% FSB. OS833 was kindly gifted from the Sweet Cordero laboratory at UCSF. OS833 was cultured in DMEM (+Glucose, +pyruvate, +GlutaMINE) GIBCO 11995-065, 10% Bovine Growth Serum, and 1% pen/strep. All cell lines were maintained in a humidified atmosphere of 5% CO_2_ at 37°C. Regular *Mycoplasma* and short tandem repeat confirmation was performed.

### Western blots

Cells were lysed with RIPA buffer (Fisher Scientific) supplemented with protease and phosphatase inhibitors (Sigma). Western blot was performed using the Bio-Rad Mini-PROTEAN Tetra System. Following transfer, nitrocellulose membranes were blocked in Odyssey Blocking Buffer (LI-COR) for 1 hour. Membranes were washed and incubated rotating overnight at 4°C with a primary antibody. Membranes were then washed in TBST and incubated with a secondary antibody (LICOR IRDye 700CW or 800CW; 1:10,000) for 1 hour. Membranes were imaged on a LI-COR Odyssey scanner.

### Cell Viability Assays

Cells were seeded at a density of 2000-5000 cells/well in 100uL medium in 96-well plates.Serial dilutions of Acla (supplied by the Neefjes Laboratory, LUMC), Doxo (Millipore Sigma, Cat #353), and Etoposide (MedChem Express, Cat#HY-13629) diluted in 100uL media were added and incubated for 72hrs. Cell viability was measured by adding 40uL Cell-Titer Glo ATP-based assay (Promega). Luminescence was read using Spectramax iD5 plate reader.

### Immunofluorescence

Cells were fixed with 4% PFA in PBS for 10 min then washed with PBS. Cells were permeabilized for 10 min with 0.3% Triton-X then blocked with 0.2% BSA in PBS for 1hr at room temperature. Cytospins were incubated with RAD51 (1:50,Sigma, Cat#ZRB1492) in antibody dilution buffer (Ventana Cat. ABD250) at 37°C in a humidified chamber followed by secondary PV Poly-HRP Anti-Rabbit IgG (Leica Microsystems Cat. PV6119) for 30 min/37°C. Alexa Fluor 568 Tyramide (Invitrogen, B40956) was then used for RAD51 detection. Slides were counterstained with DAPI (Thermo Fisher Scientific) and mounted with Prolong (Thermo Fisher Scientific), and fluorescence images were captured using a Nikon Eclipse E800 microscope (Nikon).

### In vivo cardiotoxicity experiments

All experimental procedures adhere to the *Guide for the Care and Use of Laboratory Animals* and were approved by Seattle Children’s Research Institute IACUC(18). The institute is fully AAALAC-accredited and has a Public Health Service approved Animal Welfare Assurance. C57BL/6J (7-weeks-old) male mice were purchased from Jackson Laboratory (JAX #000664) and acclimated to study for 1 week. Doxo (Millipore Sigma, Cat # 353) and Acla (provided by Neefjes Laboratory, LUMC) preparations were reconstituted with 0.9% sterile saline to a 0.5mg/mL solution. Doxo and Acla were dosed at 5 mg/kg/dose intraperitoneal injections (IP) weekly. Details of dosing schedule are found in subsequent results and figure legends. Equivalent volume of saline was delivered to vehicle mice. Mice were monitored at least three times a week for early end-point criteria which included signs of morbidity or weight loss beyond 15% of starting body weight. All mice were humanely sacrificed by carbon dioxide asphyxiation at the end of study.

### Echocardiogram

Echocardiograms (echos) were performed by a cardiologist blinded to treatment groups and during analysis; the methods of the echos are previously described(19, 20). VEVO 3100 High-Resolution Micro-Ultrasound System (VisualSonics Inc., Toronto, Canada) and MX550D transducer was used for echo measurements under isoflurane anesthesia at a 3% induction concentration and a maintenance of 1% in 100% oxygen at a flow of 1 LPM. Mice were positioned supine, and ECG leads placed for continuous monitoring during image acquisition. Heart rates were maintained between 400-600 beats per minute and heat support was provided to maintain body temperature at 37.0-37.5 °C.

Hearts were measured in the parasternal short axis at the midpapillary level of the left ventricle (LV) and taken in M-mode for end-diastole (EDD) and end-systole (ESD). Fractional shortening was calculated by [(LVEDD − LVESD) / LVEDD ∗ 100] in a parasternal short axis view for at least three heart beats. An apical 4-chamber view with pulsed doppler was used to measure early diastolic transmitral flow velocity (MV E) and late diastolic trans-mitral flow velocity (MV A) along with the myocardial performance index, calculated by [isovolumic relaxation time + isovolumic contraction time)/ aortic ejection time]. All other reported echocardiogram functional measurements were calculated in Vevo LAB (version 5.6.1, VisualSonics Inc., Toronto, Canada). Measurements were performed on five mice from each group before the start of the second exposure and 12 weeks after the final exposure, or earlier if the mouse met criteria for humane euthanasia.

### In vivo serum analysis

Prior to euthanasia, blood was collected awake by submental lancet puncture or via retroorbital bleeding under isoflurane anesthesia (2% maintenance). For complete blood counts two drops of blood were placed in EDTA coated tubes and analyzed using ElementHt5 (Heska, Loveland, CO). For additional serum analysis blood was allowed to coagulate for 20-45 minutes and spun at 2500 RPM for 15 minutes. Isolated serum was frozen at -20°C and submitted to IDEXX (Sacramento, CA).

### Tissue Preparation

Heart, lung, kidney, and liver were weighed, and tibial length measured during tissue collection. The formula (organ weight ÷ tibial length) was used to index heart weights to normalize for mouse variation in body size(21). Hearts were fixed in 10% neutral buffered formalin and paraffin embedded sections cut to 2 µm thickness in a four-chamber view. Sections were stained with hematoxylin and eosin (H&E) to be evaluated for vacuolization and trichrome special stain to highlight fibrosis. A peroxidase-based detection system (B40925, Thermofischer) and ImmPact DAB Substrate Kit (SK-4105, Vector Laboratories) was used to perform IHC with periostin primary antibody (1:100, ab215199, Abcam) and Rabbit IgG (1:100, 02-6102, Invitrogen) as a negative control. Whole slides were scanned with ZIESS Axioscan 7 (White Plains, NY) and histologic images captured using Qupath (version 0.5.1).

### Histology Analysis

A veterinary pathologist blinded to treatment groups evaluated whole heart sections and scored ventricles by severity of vacuolization, fibrosis, and immunohistochemistry stain-uptake. The right ventricle, left ventricle, and interventricular septum received a score and were summed for each heart. Scores were averaged by treatment group.

### Statistical analysis

Graphpad Prism (version 10) was used for statistical analysis. Kaplan-Meier Curves were analyzed using Log-rank (Mantel-Cox) test. Student’s t-test was used to determine significant differences between two groups and One-way ANOVA with Tukey’s Multiple Comparison for any group analysis greater than two. Statistical significance is denoted as * p-value of < 0.05, **p<0.01, *** p<0.001, **** p<0.0001 unless specified otherwise.

## Results

### Aclarubicin has equivalent *in vitro* efficacy to Doxorubicin without inducing genotoxic damage

We first compared the anti-tumor efficacy of Acla and Doxo *in vitro*. A panel of pediatric cancer cell lines were selected for which Doxo is used in upfront standard of care. Specifically, we tested CHLA10 (EwS), OS833 (OS), L-428 (HL), and Raji (BL). Etoposide was incorporated in these viability assays as a topo II poison that generates DNA breaks but does not induce chromatin damage(8). In all models, we observed nearly equivalent cytotoxicity of Acla and Doxo, and significantly decreased sensitivity to Etoposide (**Fig 1A**). We next assessed the degree of DNA damage induced by Acla, Doxo, and Etoposide at doses that induced a viability defect. CHLA10 and L-428 were exposed to 100nM Acla, 100nM Doxo, 250nM Etoposide, or vehicle for 48 hours. Protein lysates were then assessed for accumulation of DNA damage, using γ-H2AX, a marker of DNA double strand break repair. Despite these doses inducing a cytotoxic effect, only Doxo and Etoposide induced a DNA damage response (**Fig 1B**). We next assessed downstream activation of DNA damage response pathways in response to Doxo or Acla treatment through RAD51 immunofluorescence (IF), which is detected at sites of DNA damage accumulation. L-428 cells were exposed to 1μM of Acla, Doxo, or DMSO for 24hrs. While L-428 cells exhibited baseline RAD51 accumulation, Doxo-treated cells had consistently high levels, whereas Acla-treated cells showed significantly less. This suggests that RAD51 is released from chromatin by Acla, similar to the mechanism observed for histones and other proteins (**Fig 1C**). In summary, Acla shows cytotoxic efficacy comparable to Doxo, but does not induce genotoxic damage which is consistent with primary mechanism of chromatin-mediated cytotoxicity.

**Figure 1.**
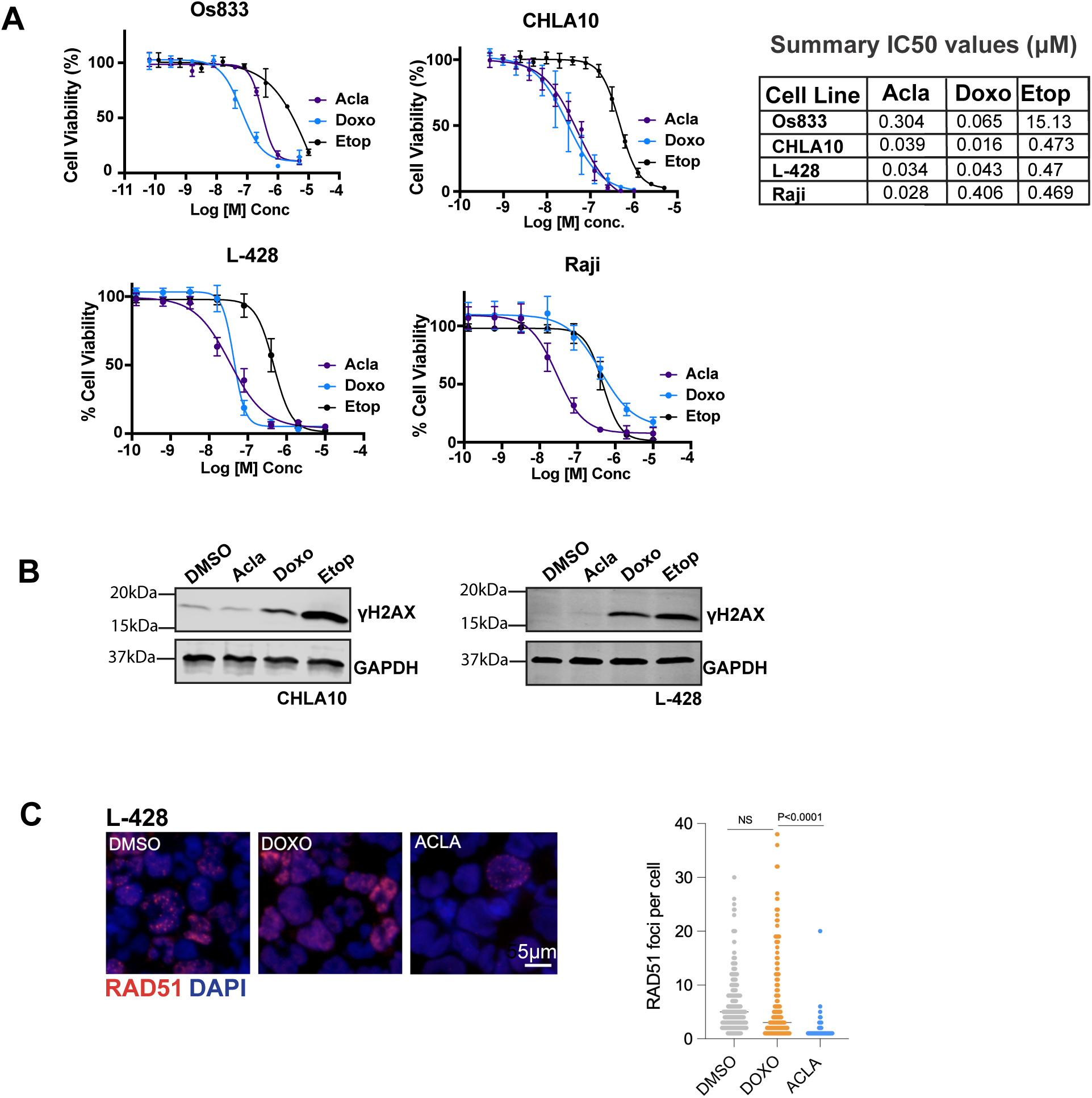
Aclarubicin has equivalent *in vitro* efficacy to Doxorubicin without inducing genotoxic damage. **A**. Dose response curves of Os833 (OS), CHLA10 (EwS), L-428 (HL), Raji (Burkitt Lymphoma). Cells were treated with serial dilutions of Aclarubicin, Doxorubicin, and Etoposide or relevant DMSO controls. Y-axis represents % viability as compared to DMSO control. Graph is representative of at least 2-3 individual experiments with 3 technical replicates. Table shows IC50 values**. B.** Immunoblot of phospho-ψH2AX protein expression (representative image of n=2) of CHLA10 (EwS) and L-428 (HL) cells treated with 100nM Aclarubicin, 100nM Doxorubicin, or 250nM Etoposide for 48hrs. **C.** DAPI and RAD51 staining of L-428 cells treated with DMSO, 100nM Doxorubicin, or 100nM Aclarubicin for 48hrs (representation image of n=2). Quantification of RAD51 foci nuclear staining per cell. Statistical significance calculated using student t-test (* p<0.05, ** p<0.01 ***p<0.001)

### Modeling Doxorubicin Cardiotoxicity *in vivo*

There are multiple published mouse models for evaluating Doxo induced cardiotoxicity using different mouse strains and dosing. Thus, we first sought to validate induction of Doxo-induced cardiotoxicity in the C57BL/6J strain. Recent studies inducing cardiotoxicity use a cumulative Doxo dose of 20-25 mg/kg, and evaluate for cardiac changes by histology 1-13 weeks after the last dose of Doxo is delivered(22–24). Given this, we exposed non-tumor bearing C57BL/6J mice to saline or 5mg/kg Doxo weekly for total cumulative doses of 20mg/kg, 25mg/kg, or 30mg/kg. Mice were observed for up to 13 weeks after the last dose of Doxo, at which point all animals were euthanized (**Fig 2A**).

**Figure 2.**
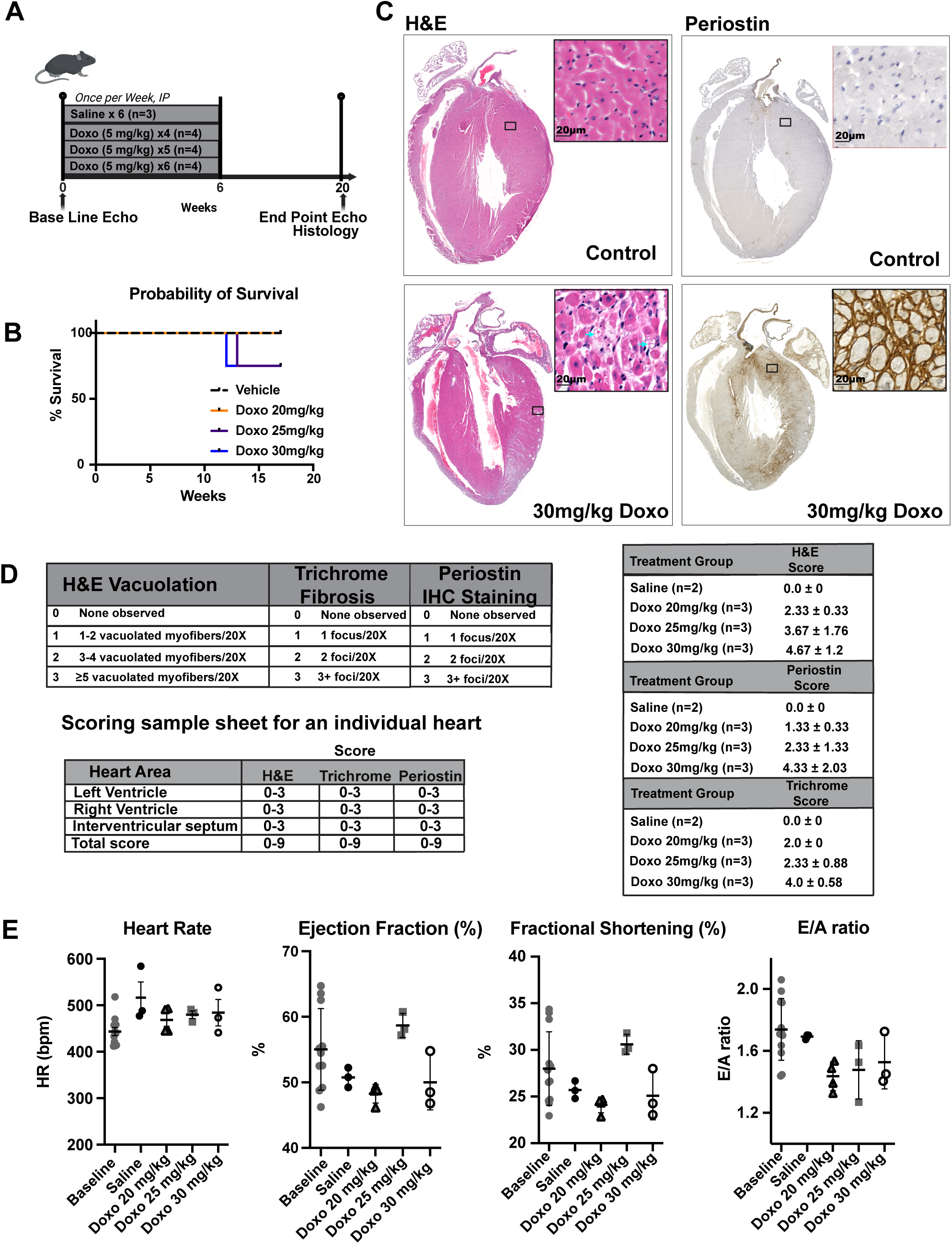
Modeling Doxorubicin Cardiotoxicity *in vivo.* **A**. Schema of xenograft study in C57BL/6J mice treated with dose escalation of Doxorubicin. **B.** Kaplan-Meier overall survival curves of individual treatment groups. **C**. H&E and Periostin staining of whole heart scans with magnification of blue boxed region. Blue arrow indicates areas of vacuolization. Representative images are from animals treated with total cumulative 30mg/kg or saline. **D.** Histology scoring system for H&E, Trichrome, and Periostin and sample scoring sheet for an individual heart. Histology scores from H&E, trichrome, and periostin by treatment groups (Mean ± SD). There was no statistical significance between treatment groups. **E.** Scatter plots of echocardiogram measurements of the left ventricle and mitral valve at baseline, saline, and treatment groups (Mean ± SEM). Reference ranges: HR -400-550 beats per minute; EF- 47%-70%; FS 24%-40% No statistical significance between groups.

Echocardiograms were performed at baseline and at time of sacrifice (**Fig 2A**). Only two mice, one that received a total cumulative dose of 25mg/kg, and one that received 30mg/kg, required humane euthanasia prior to endpoint due to weight loss >15% (**Fig 2B**). All mice that received Doxo had histologic evidence of cardiotoxicity demonstrated by increased vacuolation on H&E, and marked fibrotic changes throughout the heart, with increased periostin (**Fig 2C**) and trichrome staining (data not shown). Although not statistically significant due to sample size, histopathologic scoring of H&E vacuolation, trichrome fibrosis, and periostin IHC staining suggested a dose dependent increase in cardiotoxicity (**Fig 2D**).

As echocardiogram is the current gold standard for detecting anthracycline induced cardiotoxicity in pediatric patients, we aimed to correlate our histologic findings with mouse echocardiograms. Specifically, we looked at left ventricular Ejection Fraction (EF) and Fractional Shortening (FS), markers used currently to clinically diagnose anthracycline cardiotoxicity(4).

Additionally, we measured E/A ratio, which is used to detect left ventricular diastolic dysfunction. Overall, we observed a variable range of echo results. No statisctical significance was identified, however there was a trend towards a reduced EF, FS, and E/A ratio in mice receiving either 20mg/kg or 30mg/kg of Doxo (**Fig 2E**). In conclusion, doses of 20mg/kg-30mg/kg Doxo reproducibly induce histologic evidence of cardiotoxicity in a C57BL/6J mouse model; but there were no reproducible functional changes by echo.

### Aclarubicin is safe to deliver after Doxorubicin

After confirming that 20–30 mg/kg Doxo induces cardiotoxicity, we tested the safety of Acla following Doxo exposure to model relapse treatment, hypothesizing improved tolerance due to minimal genotoxic stress. Mice were treated with 5mg/kg Doxo weekly for a total of 4 doses (total cumulative 20mg/kg). After a 4 week rest period, mice were randomly assigned to receive four additional doses of 5mg/kg Doxo (Doxo-Doxo), 5mg/kg Acla (Doxo-Acla), or Saline (Doxo-Saline) weekly (additional total cumulative dose of 20mg/kg). Mice were observed for up to 12 weeks after completion of the second course of therapy, or humanely euthanized if they exhibited >15% weight loss or other significant toxicities (**Fig 3A**). As expected, the second course of Doxo was found to be highly toxic to the mice. Only 20% of mice were alive at study endpoint (3/15) and the group showed a median survival of 12 weeks from first dose of Doxo. This is in comparison to the Doxo-Saline group, where 50% of the mice were alive at study endpoint (5/10), with a median survival of 20.5 weeks. Strikingly, the Doxo-Acla group showed no evidence of additional toxicity and 73% of mice were alive (11/15) at study endpoint, with an undefined median survival. While the survival of the Doxo-Saline group relative to Doxo-Doxo was superior, it just missed the threshold for statistical significance (p=0.070). However, as compared to the Doxo-Acla group, there was a significantly higher rate of toxic mortality in the Doxo-Doxo group (p=0.0098) (**Fig 3B**). There was no statistical significance in the rate of toxic mortality between the Doxo-Acla group and the Doxo-Saline group (p=0.33).

**Figure 3.**
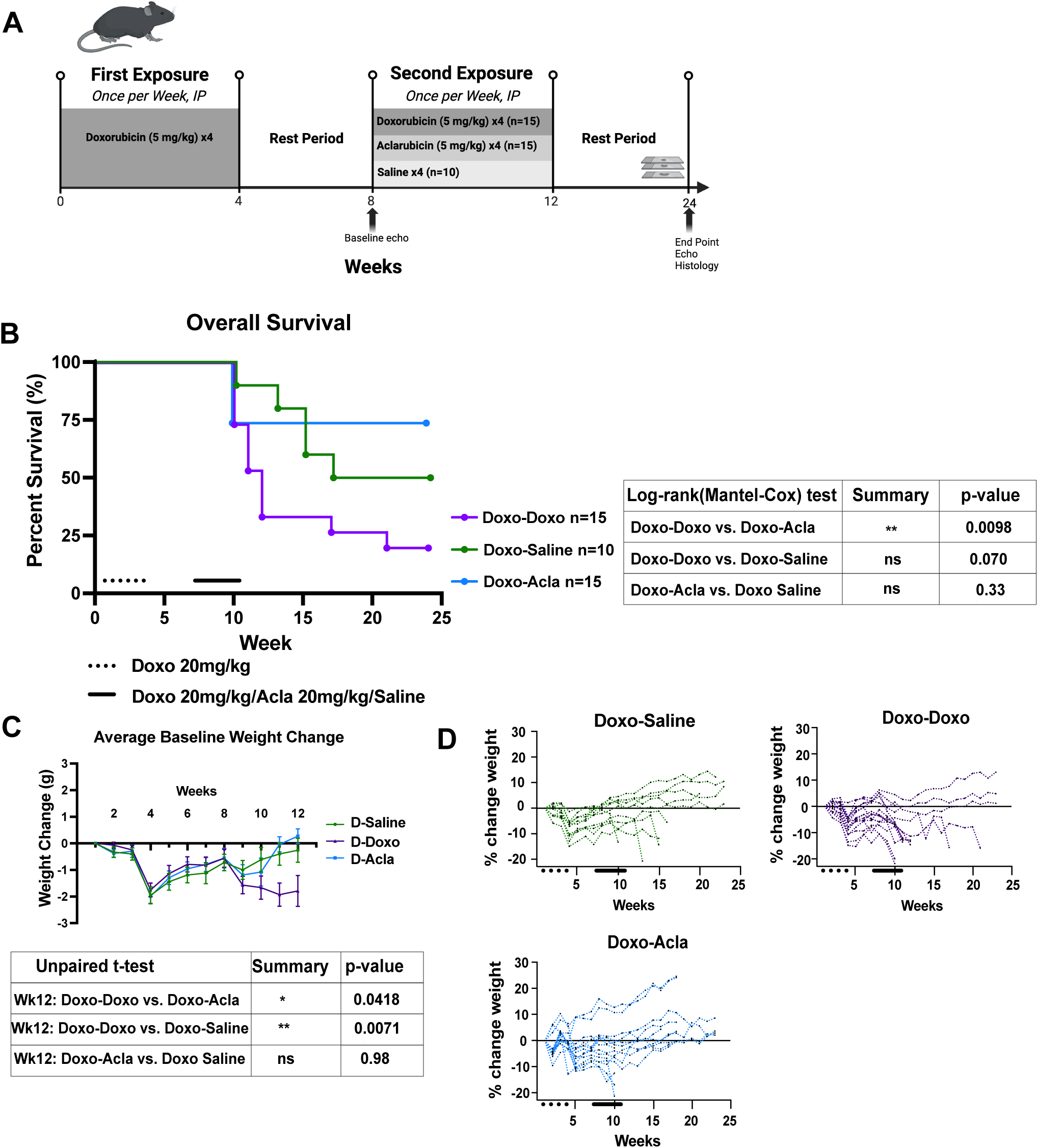
Aclarubicin is safe to deliver after Doxorubicin. **A**. Schema of xenograft study in C57BL/6J mice treated with 20mg/kg of Aclarubicin, 20mg/kg Doxorubicin, or Saline after initial 20mg/kg Doxorubicin treatment. **B.** Kaplan-Meier overall survival curves of individual treatment groups demonstrate a statistically significant increased toxic mortality in only mice that receive a second course of Doxorubicin. Statistical significance measured by Log-rank test, with table of p-values. **C.** Change in weight (g) from baseline of each individual treatment group from weeks 1-12. The Doxo-Doxo treatment group had a statistically significant weight loss as compared to Doxo-Acla or Doxo-Saline group at week 12, at the completion of the second course of treatment. Statistical significance calculated using multiple student t-test (* p<0.05, ** p<0.01 ***p<0.001). **D.** Individual percentage change in weight from baseline until endpoint.

Striking changes to baseline weights further reflect the significant differences in toxic morbidity. All mice lost weight after the first Doxo dose (Week 4) but recovered by Week 8, prior to the second course of treatment (**Fig 3C**). However, only the Doxo-Doxo group experienced statistically significant severe weight loss during and after the second course of therapy (**Fig 3C-D**). The few surviving mice in the Doxo-Doxo group largely failed to gain weight, which was not observed in the Doxo-Acla or Doxo-Saline groups.. These findings demonstrate that Acla is safe following Doxo and does not increase toxic mortality.

### Accumulation of significant cardiac toxicity is not observed with Aclarubicin

Given the striking increase in toxic mortality seen in the Doxo-Doxo group, we hypothesized this was secondary to cardiotoxicity. We first evaluated for histologic changes. Representative H&E, and Trichrome images from one mouse from each treatment group are shown in **Fig 4A**. Histopathologic scoring of H&E vacuolation (Doxo-Saline, n=4; Doxo-Doxo, n=9; Doxo-Acla, n=3) and trichrome (Doxo-Saline, n=9; Doxo-Doxo, n=13; Doxo-Acla, n=13) fibrosis were performed in a subset of mice (**Fig 4B**). Surprisingly, while we observed mild vacuolation, and mild fibrosis (evident by increased trichrome staining) in all groups, this was not increased in the Doxo-Doxo group (**Fig 4C**).

**Figure 4.**
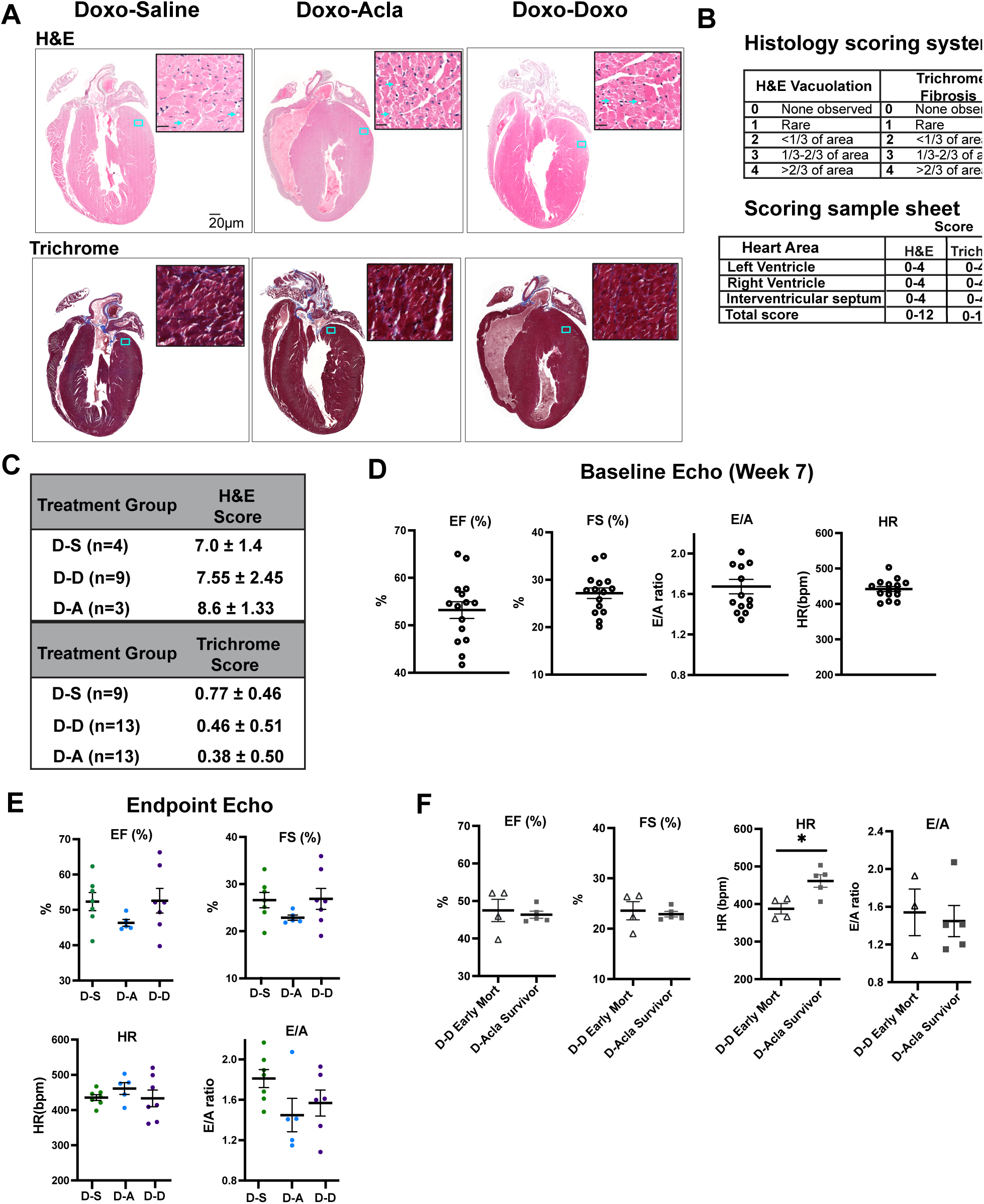
Accumulation of significant cardiac toxicity is not observed with Aclarubicin. **A**. H&E and trichrome staining of whole heart scans with magnification of blue boxed region. Blue arrow indicates areas of vacuolization. Representative images are from animals at endpoint in each individual treatment group. **B**. Histology scoring system for H&E and trichrome, and sample scoring sheet for an individual heart. **C.** Histology scores from H&E and trichrome by treatment groups at endpoint (Mean ± SD). There was no statistical significance between treatment groups. Scatter plots of echocardiogram measurements of the left ventricle and mitral valve at **D.** baseline (week 7) prior to mice receiving a second course of anthracyclines or saline, **E.** at study endpoint or time of humane euthanasia, and **F.** between Doxo-Doxo animals that experienced early mortality and Doxo-Acla mice that reached study endpoint. Reference range-HR -400-550 beats per minute; EF- 47%-70%; FS 24%-40%. Mean ± SEM is plotted, and statistical significance was calculated using student t-test which did not show any statistical significance between groups.

Next, we performed echos on nineteen mice (Doxo-Doxo n=7, Doxo-Saline n=7, Doxo-Acla n=5) that were randomly selected prior to initiation of the second course of therapy. Baseline cardiac function was performed at week 7 prior to the second course of anthracyclines. Baseline echos showed high inter-mouse variability in all cardiac measurements-EF, FS, HR, and E/A (**Fig 4D**). Treatment endpoint echos showed no significant changes were evident across treatment groups (**Fig 4E**). All mice followed by echocardiogram in the Doxo-Acla group reached study endpoint, as compared to only 3/7 mice in the Doxo-Doxo group. Echos of surviving mice in the Doxo-Acla group and early mortality Doxo-Doxo group failed to identify significant differences (**Fig 4F**). Thus, despite a significantly higher rate of early toxic mortality in the Doxo-Doxo group there was no definite histologic or echocardiographic evidence of augmented cardiotoxicity.

### End organ dysfunction is not observed with Aclarubicin delivery

To define other potential causes of early mortality in Doxo-Doxo mice relative to Doxo-Acla mice, we evaluated for additional evidence of organ dysfunction, specifically bone marrow, kidney, and liver. Complete blood counts (CBC) was obtained at time of euthanasia, and we specifically analyzed the differences between Doxo-Acla mice reached study endpoint versus Doxo-Doxo mice that experienced early mortality. Doxo-Doxo mice that experienced early mortality had significantly lower total white blood cells, specifically lymphocytes. No significant differences in platelets or hemoglobin were observed (**Fig 5A**). Comprehensive metabolic profiles were obtained from mice that reached study endpoint from all three groups. Doxo-Doxo survivors had lower total calcium and phosphorous than Doxo-Acla survivors, and no differences in potassium levels were observed (**Fig 5B**). Doxo-Doxo survivors had higher ALT compared to Doxo-Saline survivors, but there was no difference in total bilirubin (**Fig 5C**). Finally, as measured by BUN and creatinine, no differences in kidney function were observed (**Fig 5D**). Thus, no evidence of end organ damage was evident in Doxo-Acla treated mice despite a cumulative anthracycline dose of 40 mg/kg. In summary, no end-organ damage was observed in Doxo-Acla mice despite a cumulative anthracycline dose of 40 mg/kg. In contrast, Doxo-Doxo mice with the same dose experienced significant early mortality, though the cause was unclear at necropsy.

**Figure 5.**
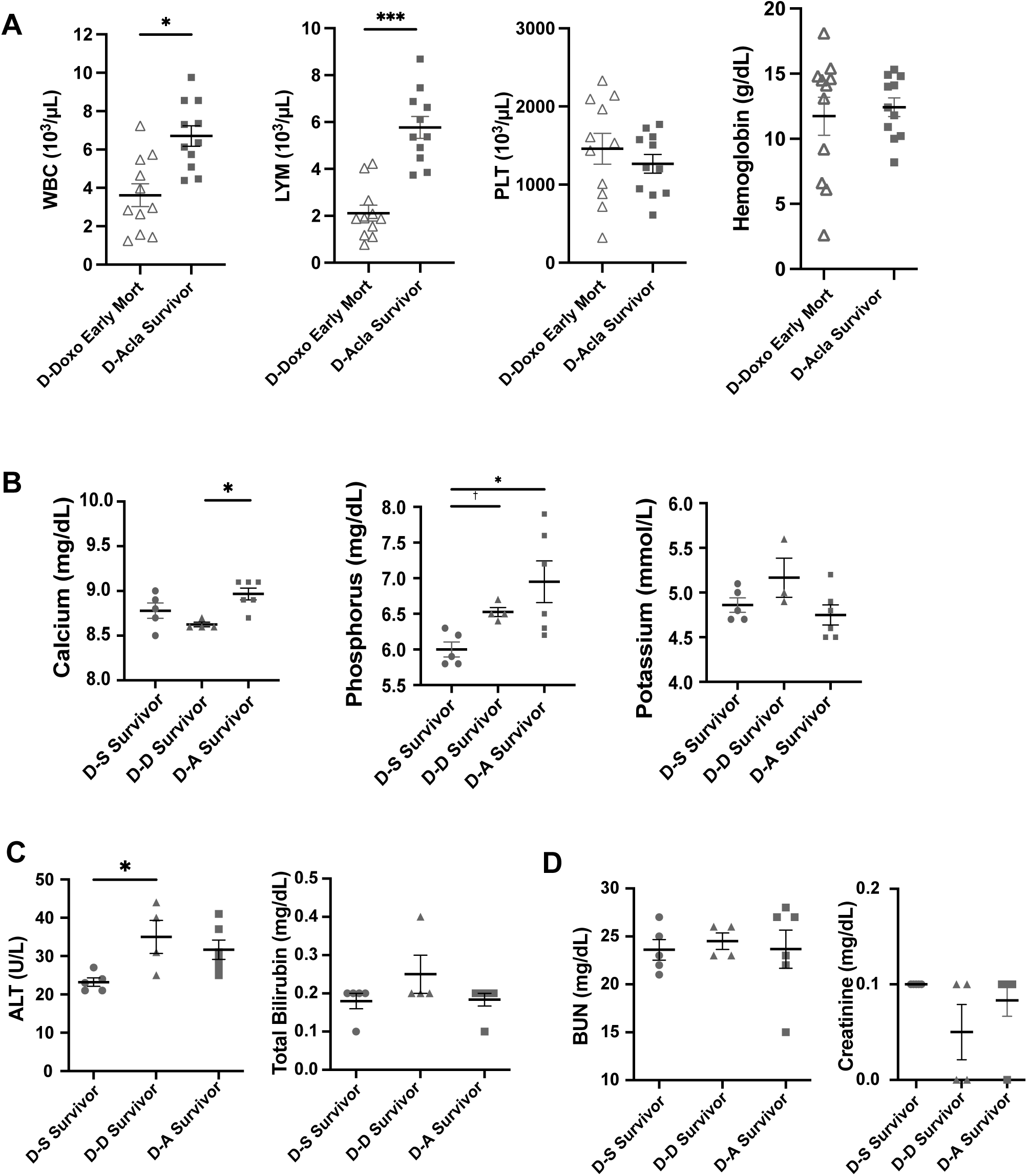
End organ dysfunction is not observed with Aclarubicin delivery. **A**. Scatter plots of complete blood count measurements of Doxo-Doxo mice that experienced early mortality and Doxo-Acla mice that reached study endpoint. Reference range: WBC: 3.5 x10^3^/uL-6.9 x x10^3^/uL; Lymphocytes: 2.92 x10^3^/uL-5.86 x x10^3^/uL; Platelets 727x10^3^/uL-1177 x10^3^/uL; Hemoglobin: 10.7g/dL-12.3g/dL. Scatter plots of **B.** electrolytes, **C.** liver function tests and **D.** kidney function analysis of each individual treatment group of animals that reached study endpoint and did not experience early mortality. Calcium: 8.8 mg/dL-9.2 mg/dL; Phosphorous 6.1 mg/dL-7.5 mg/dL; Potassium: 3.8 mmol/L-4.2 mmol/L; ALT 18 U/L -94U/L; Total bilirubin 0.3mg/dL-0.7mg/dL; BUN 21.8 mg/dL-32.38 mg/dL; Creatinine 0.1mg/dL-0.15mg/dL. Mean ± SEM plotted, and statistical significance calculated using multiple student t-test.

## Discussion

Anthracyclines, one of the most important class of chemotherapy agents, is limited by cumulative dose-dependent toxicities. Acla, which induces chromatin damage without significant DNA damage, offers a promising alternative(8–11). Our data demonstrate that Acla exhibits anti-tumor efficacy comparable to Doxo across multiple pediatric *in vitro* models, without inducing genotoxic damage. Further, *in vivo* models recapitulating delivery of anthracyclines after Doxo exposure, demonstrate the superior safety profile of Acla. *In vivo*, Acla administered after prior Doxo exposure did not exacerbate toxicity, unlike additional Doxo, which led to increased toxic mortality.

Pediatric malignancies are characterized by widespread epigenetic dysregulation, with the highest rate of mutations occurring in epigenetic regulators(25) (26–28). Malignancies without these mutations, still demonstrate widespread epigenetic disruption, such as EwS(29, 30). We speculate that the superior efficacy of anthracyclines in pediatric cancers may indeed stem from their ability to disrupt epigenetic programs. Indeed, a small case series from Japan demonstrated activity of cisplatin and Acla in intracranial alveolar rhabdomyosarcoma(31). Our studies, which show equal efficacy of Acla to Doxo, and superior efficacy to etoposide, which solely induces DNA damage, further support the activity of Acla in pediatric malignancies. Further validation of Acla’s efficacy in mouse xenograft studies are ongoing.

Survival rates for pediatric malignancies have drastically improved. However, treatment-related toxicities, linked to anthracycline exposure, ie cardiovascular disease and secondary malignancies, including breast cancer independent of radiation(32), are increasing in survivors(33, 34) (4, 35). Our studies highlight the need to develop preclinical models that reproducibly model anthracycline-induced toxicities. Previous reports demonstrate a wide range of time for detection of chronic cardiac changes, anywhere between 4- and 13-weeks post-exposure(22–24, 36). Our findings support that histologic evidence of chronic cardiac changes take time to develop. As we initially hypothesized that a second course of Doxo would exacerbate cardiac changes, we were surprised that despite increased toxic mortality in the Doxo-Doxo group (**Fig. 3**), cardiotoxicity was not the cause. This was likely due to the significantly shortened median survival for the Doxo-Doxo group, only 12 weeks, or two weeks after the final dose. As our initial findings detected cardiac remodeling 12 weeks after the final dose of Doxo (**Fig. 2**), results from both models presented strongly support that histologic cardiac remodeling changes take greater than 8-10 weeks to occur.

Importantly, our results validate that high doses of Doxo are unsafe, and the superior survival of the Doxo-Acla group underscores the safety of Acla delivery after Doxo exposure. Given median time of survival for Doxo-Doxo mice, our model suggests that acute rather than chronic toxicity was captured. One mechanism of acute anthracycline toxicity is driven by systemic inflammation(37). Doxo has been shown to activate the innate immune system, in particular Toll-like receptor 4 (TLR4), JAK/STAT signaling pathway, and inflammatory cytokines leading to multiorgan damage and potentiating cardiac changes(37–39). Interestingly, recent preclinical studies report that Acla can suppress these same inflammatory pathways(40). Our observation of improved, but not statistically significant, survival in the Doxo-Acla group as compared to the Doxo-Saline group suggests that Acla may even exert a protective effect, potentially by mitigating inflammatory responses. Validation of this differential inflammatory response will be important to demonstrate in future experiments.

In summary, our results demonstrate that Acla exhibits potent anti-tumor activity across multiple pediatric malignancies with a superior safety profile, allowing its administration after prior Doxo exposure. This highlights the significance of chromatin damage as a key mechanism of action for anthracyclines and supports the need for further investigation of Acla in pediatric cancer therapy.

## Conflict of Interest Statement

Dr. Jacques Neefjes is a shareholder in NIHM that aims to produce Aclarubicin for clinical use. The other authors have no competing interests to disclose.

## Acknowledgments

The authors thank members of the Lawlor, Sarthy, Neefjes, Olson, and Maves labs for helpful discussion, and the staff at the Seattle Children’s Research Institute histology and microscopy core and animal core. We would also like to thank colleagues at University of Washington, Department of Veterinary Medicine. Grant and gift support for this work is gratefully acknowledged and was provided by the following sources: NIH/NCI 5K12CA076930 (SSG), NIH/NCI Loan Repayment Grant 1L40CA264716 (SSG), CURE Childhood Cancer (SSG, ERL, JFS), Sam Day Foundation (LM, ERL, JFS), Burroughs Wellcome Fund (JFS), Andy Hill CARE Fund (JFS), Sunbeam Foundation (JFS), NIH/NCI 5K08CA256167 (JFS), NIH/NCI 1R21CA280139 (LM, ERL, AO).

## Abbreviation Table

Doxo: Doxorubicin
Acla: Aclarubicin
EwS: Ewing sarcoma
OS: Osteosarcoma
HL: Hodgkin Lymphoma
BL: Burkitt Lymphoma
IF: Immunofluorescence
IHC: Immunohistochemistry
IP: Intraperitoneal
Doxo-Saline: Doxorubicin-Saline treatment group
D-S: Doxorubicin-Saline treatment group
Doxo-Acla: Doxorubicin-Aclarubicin treatment group
D-A: Doxorubicin-Aclarubicin treatment group
Doxo-Doxo: Doxorubicin-Doxorubicin treatment group
D-D: Doxorubicin-Doxorubicin treatment group
LV: Left ventricle
Echo: Echocardiogram
HR: Heart rate
FS: Fractional shortening
EF: Ejection fraction
AYA: Adolescent and young adult
Topo II: Topoisomerase II
dsDNA: DNA damage double strand break
CBC: Complete blood count
ALT: Alanine Aminotransferase
AST: Aspartate aminotransferase
LYM: Total lymphocyte count
PLT: Total Platelet count
TLR-4: Toll-like receptor 4

